# Reprogramming a high robust *Geobacillus thermoglucosidasius* for efficient synthesis of polymer-grade lactic acid under extremely high temperature (60°C)

**DOI:** 10.1101/2023.04.14.536835

**Authors:** Jiongqin Liu, Xiao Han, Fei Tao, Ping Xu

## Abstract

Optically pure lactic acid is an important precursor for the synthesis of biodegradable plastic polylactic acid. *Geobacillus thermoglucosidasius* is a thermophilic bacterial chassis with industrial potential. In this study, we reprogrammed this chassis to efficiently produce polymer-grade lactic acid by combining rational and semi-rational design strategies. We obtained optically pure L-lactic acid- and D-lactic acid-producing strains, GTD17 and GTD7, respectively, via rational strategies: constructing a lactic acid synthesis module, deleting by-product synthesis, and enhancing lactic acid production. At an extremely high temperature (60°C), the engineered strain GTD17 produced 94.2 g L^−1^ of L-lactic acid with an overall yield and productivity of 91.5% and 2.0 g L^−1^ h^−1^, respectively, and the optical purity of L-lactic acid was 99.5%. We then performed semi-rational adaptive evolution on the engineered strain GTD7 to obtain an optically pure D-lactic acid-producing strain GTD7-144; at the extremely high temperature (60°C), it produced 153.1 g L^−1^ of D-lactic acid with an overall yield and productivity of 93.0% and 3.2 g L^−1^ h^−1^, respectively. The optical purity of the D-lactic acid was 99.6%. Genome resequencing and analysis of the strain GTD7-144 revealed potential genetic targets that may improve production performance. This study represents an advancement in the high-temperature production of polymer-grade lactic acid by *G. thermoglucosidasius*.

## Introduction

Lactic acid is an important, naturally occurring, small-molecule bulk chemical. Based on chirality, it can be divided into L-lactic acid and D-lactic acid, which are produced by the conversion of pyruvate, catalyzed by L-lactate dehydrogenase and D-lactate dehydrogenase, respectively^1^. Optically pure L-lactic acid and D-lactic acid can be used as precursors to produce biodegradable plastic polylactic acids (PLA, which includes PLLA and PDLA)^2–5^. PLA has received increasing attention in many fields due to its biodegradability and biocompatibility^6^. Changing the ratio of the two chiral precursors in PLA can improve its thermal resistance and mechanical properties^7–12^. As an alternative to non-biodegradable plastics, the application of PLA can alleviate the problem of plastic pollution, which conforms to the concept of green environmental protection and is beneficial to the ecological balance of the Earth^6^. In 2019, the worldwide plastic production was 368 million tons^13^. The resulting plastic pollution has prompted market demand for biodegradable plastics. Therefore, the low-cost and efficient production of optically pure L-lactic acid and D-lactic acid is urgently needed, and their market demand will be very high.

*Geobacillus thermoglucosidasius* is a thermophilic, gram-positive, aerobic, or facultative anaerobic bacterial chassis with industrial potential^14, 15^. *G. thermoglucosidasius* can grow rapidly between 40 to 70°C, and the optimal growth temperature is around 60°C, at which its growth level is comparable to that of *Escherichia coli*^16^. *G. thermoglucosidasius* has a very wide spectrum of substrate utilization and catabolic versatility, and can metabolize a wide range of C_5_ and C_6_ sugar monomers and oligomers^17^. In addition, researchers have developed several genetic manipulation methods that are suitable for this strain to facilitate its metabolic engineering development^18–23^. Therefore, using *G. thermoglucosidasius* as a chassis for fermentation production at high temperatures has practice prospects, which has the following advantages. This can reduce the cooling cost of large-scale exothermic fermentation, which becomes increasingly obvious with the increase in fermentation scale^24–26^. This reduces the risk of contamination by mesophilic microorganisms that cannot survive at high temperatures^27^. It is beneficial to the industrial biological process of simultaneous saccharification and fermentation and can reduce the cost of enzyme use because the optimal reaction temperature of the enzymes used in this process is relatively high^28–30^. These advantages contribute to the reduction in production costs. In addition, it can make unfavorable reactions thermodynamically feasible in mesophilic microorganisms^31^. It can provide fermentation media with good properties, such as lower viscosity, faster substrate diffusion rates, and better substrate solubility^20^. It is beneficial for the production, collection, and extraction of volatile products, as it can reduce the effects of product inhibition and product toxicity^32^. These advantages are conducive to improving the production efficiency. All these advantages of high-temperature fermentation are very important for the production of cost-sensitive small-molecule bulk chemicals. Therefore, in this study, we focused on reprogramming this chassis to efficiently produce optically pure L- and D-lactic acids at high temperatures.

Adaptive evolution is a semi-rational method used in laboratories to improve the fermentation performance of strains and can create infinite possibilities^33^. Adaptive evolution has been used by researchers to improve the utilization rate of substrates by strains^34–36^, enhance the resistance of strains to toxic substrates and products^37–39^, and improve the adaptability of strains to extreme environments such as low pH, high osmotic pressure, and high temperature^40–42^. Researchers may obtain desired super strains to adapt to these conditions and produce the required compounds with high efficiency through adaptive evolution.

In this study, using *G. thermoglucosidasius* DSM 2542 as the initial strain, we obtained the optically pure D-lactic acid-producing strain GTD7 through the following series of rational metabolic engineering strategies: construction of a D-lactic acid synthesis module, deletion of by-product synthesis, and enhancement of D-lactic acid production. Using the engineered strain GTD7 as the initial strain, we obtained the optically pure L-lactic acid-producing strain GTD17 through the construction and enhancement of L-lactic acid synthesis and deletion of by-product synthesis. Finally, we performed semi-rational adaptive evolution of the engineered strain GTD7 to obtain the optically pure D-lactic acid-producing strain GTD7-144, which exhibited excellent D-lactic acid production performance (Fig. 1). Potential genetic targets were identified using genome resequencing and analysis of strain GTD7-144. In addition, we explored solutions for programmed cell death and lysis in *G. thermoglucosidasius*.

## Experimental procedures

### Bacterial strains and plasmids

All bacterial strains and plasmids used in this study are listed in Table 1 and Table S1, respectively. *Escherichia coli* DH5α strain was used for the plasmid construction. *G. thermoglucosidasius* DSM 2542 was used as the wild-type parent strain for the construction of the engineered strains. *Bacillus licheniformis* BJQX and *Bacillus coagulans* H-2 were used to amplify the corresponding D-lactate dehydrogenase and L-lactate dehydrogenase, respectively^4^. *B. coagulans* H-2 was used to amplify the *pfk* and *pyk* genes. The *E. coli*-*G. thermoglucosidasius* shuttle plasmid pUB-sfGFP was used for markerless gene deletion and integration in engineered strains^22^.

### Media and culture conditions

1. *E. coli* was cultured in Luria-Bertani (LB) broth (10 g L^−1^ NaCl, 5 g L^−1^ yeast extract, 10 g L^−1^ tryptone) or on LB agar (10 g L^−1^ NaCl, 5 g L^−1^ yeast extract, 10 g L^−1^ tryptone, 1.8 g L^−1^ Agar) at 37°C on a rotary shaker (200 rpm) supplemented with ampicillin (100 μg mL^−1^) when required. The seed of *G. thermoglucosidasius* was grown in TSB medium at 60°C on a rotary shaker (200 rpm). The TSB medium contains 15 g L^−1^ tryptone, 5 g L^−1^ soya peptone and 5 g L^−1^ NaCl. TSA medium is similar to TSB medium and additionally contains 1.8 g L^−1^ Agar. When necessary, kanamycin (12.5 μg mL^−1^) was added to the TSB and TSA media. Fermentation medium that was a modified ASYE mineral salt medium named GTP medium was used for anaerobic fermentation of *G. thermoglucosidasius* at 60°C^22^. GTP medium contains 5 g L^−1^ yeast extract, 3 g L^−1^ Na_2_HPO_4_, 3 g L^−1^ KH_2_PO_4_, 1 g L^−1^ NH_4_Cl, 0.48 g L^−1^ MgSO_4_, 0.5 g L^−1^ NaCl, 0.42 g L^−1^ citric acid, 0.028 g L^−1^ FeSO_4_·7H_2_O, 0.01 g L^−1^ thiamin, Trace Metal Mix (4.4 mg L^−1^ NiSO_4_·6H_2_O, 2.86 mg L^−1^ H_3_BO_3_, 1.81 mg L^−1^ MnCl_2_·4H_2_O, 0.39 mg L^−1^ Na_2_MoO_4_·2H_2_O, 0.222 mg L^−1^ ZnSO_4_·7H_2_O, 0.079 mg L^−1^ CuSO_4_·5H_2_O, 0.049 mg L^−1^ Co(NO_3_)_2_·6H_2_O), 3.1 mg L^−1^ biotin, and 0.1 g L^−1^ betaine. Glucose or xylose was used as carbon source for fermentation. The electroporation medium contains 0.5 M mannitol, 0.5 M sorbitol, and 10% (v/v) glycerol. The 2TY medium for electroporation incubation contains 16 g L^−1^ tryptone, 10 g L^−1^ yeast extract, and 5 g L^−1^ NaCl. 2TYA medium is similar to 2TY medium and additionally contains 1.8 g L^−1^ Agar.

### Plasmid and strain construction

All DNA manipulation and general molecular biology techniques were conducted according to the standard protocols. All primers used in this study are listed in Table S2. The linearized vector was obtained by double digestion of the pUB-sfGFP plasmid with *Bam*HI and *Hind*III. The primer pairs LDH-Up-F/LDH-Up-R and LDH-Down-F/LDH-Down-R were used to amplify the upstream and downstream homology arms by using polymerase chain reaction (PCR) technology from the genome of *G. thermoglucosidasius*, respectively. The primers LDH-DLDH-F and LDH-DLDH-R were used to amplify the *D-ldh* gene by using PCR technology from the genome of *B. licheniformis* BJQX. The primers LDH-Up-F and LDH-Down-R were used to amplify the LDH-DLDH-UD fragment by using splicing by overlap extension polymerase chain reaction (SOE-PCR)^43^ from the templates of upstream homology arm, *D-ldh* gene, and downstream homology arm. The linearized pUB-sfGFP vector and the LDH-DLDH-UD fragment were used to obtain the pUB1 plasmid by using seamless cloning with the ClonExpress Ultra One Step Cloning Kit (Vazyme, Nanjing, China). Primers pUBTY-F and pUBTY-R were used to verify the sequence of the fragments which inserted into the MCS region of pUB-sfGFP. Recombinant plasmids required for other genetic manipulations were obtained using the similar process that described above (Table S1).

The genetic manipulations of *G. thermoglucosidasius* were performed as described by Yang^22^ *et al.* Briefly, the recombinant plasmids were introduced into *G. thermoglucosidasius* by electroporation (25 kV/cm, 10 μF, 600 Ω). The resulting transformants were cultured in 2TY medium and then cultured on 2TYA plates containing kanamycin at 68°C for 12 h to generate single crossover strains. Then they were cultured in TSB medium at 60°C for 10 h and passaged for 2 –5 generations. The double crossover strains were then screened on TSA plates using green fluorescence. Finally, the corresponding primers were used for PCR verification to obtain the correct engineered strains (Table 1). The DNA polymerase was purchased from Takara Biochemicals (Shanghai, China). The restriction enzymes were purchased from New England Biolabs (Beverly, MA). The DNA fragments were recovered using the Agarose Gel DNA Column Recovery Kit (GENERAY, Shanghai, China).

The recombinant plasmids were extracted using the TIANprep Rapid Mini Plasmid Kit (TIANGEN, Beijing, China). Genomic DNAs of bacteria were extracted using the Wizard Genomic DNA Purification Kit (Promega, Madison, WI, USA).

### Fermentation production of L-lactic acid and D-lactic acid

The engineered strains of *G. thermoglucosidasius* were first cultured in a 50 mL culture shake flask containing 30 mL of TSB medium with a rotation rate of 200 rpm at 60°C for 12 h to make them grow stronger, and the inoculation volume was 0.3% (v/v). Then they were transferred to a 500 mL culture shake flask containing 150 mL of TSB medium with a rotation rate of 200 rpm at 60°C for 12 h which were used as the seed culture for fermentation, and the inoculation volume was 0.3% (v/v). The seed culture used for fermentation were inoculated into a 5 L fermentation bioreactor (Bailun Bio, Shanghai, China) containing 2.7 L of fresh GTP medium, and the inoculation volume was 10% (v/v), so that the final initial fermentation liquid volume was 3 L. The culture pH was automatically maintained at 7.0 by adding 25% (w/v) Ca(OH)_2_. The cultivation was carried out at 60°C and 80 rpm. In the fermentation process, glucose (or xylose) was added to the bioreactor to maintain a suitable sugar concentration when the sugar concentration was between 0 and 40 g L^−1^, and the fermentation time was controlled within 50 h. The initial glucose concentration was 40 g L^−1^, 80 g L^−1^ or 100 g L^−1^ depending on different engineering strains, and the initial xylose concentration was 40 g L^−1^. Samples were collected periodically to determine the cell density and the concentrations of glucose (or xylose), L-lactic acid, D-lactic acid and other by-products. The verification of the fermentation performances of the engineered strains in this study were all carried out in a 5 L fermentation bioreactor.

### Analytical techniques

The concentrations of glucose were quantified by an SBA-40D biosensor analyzer (Institute of Biology, Shandong Academy of Sciences, China). The cell density (OD_600_) was measured by a spectrophotometer (V-1200 of Mapada, Shanghai, China). The concentrations of L-lactic acid, D-lactic acid, formic acid, acetic acid, ethanol, succinic acid, and xylose were measured via a high-performance liquid chromatography (HPLC) system (Agilent 1260 series, Hewlett-Packard, USA) equipped with a Bio-Rad Aminex HPX-87H column (300 × 7.8 mm) and a differential refractive index detector. The working conditions were as follows: 5 mM H_2_SO_4_ as the mobile phase with a flow rate of 0.5 mL min^−1^ at 55°C and th e injection volume was 10 μL^44, 45^. The optical purity of L-lactic acid and D-lactic acid were measured via a HPLC system (Agilent 1260 series, Hewlett-Packard, USA) equipped with a SCAS Sumichiral OA-5000 column (150 × 4.6 mm) and a diode array detector. The UV absorption wavelength was set to 254 nm. The working conditions were as follows: 2 mM CuSO_4_ as the mobile phase with a flow rate of 0.8 mL min^−1^ at 30°C and the injection volume was 10 μL. The samples were processed as described by Han^3^ *et al.* Briefly, the samples were heated at 100°C for 10 min, then 2 mL was taken into a 1 00 mL volumetric flask and acidified with 2 mL 2 M H_2_SO_4_ for 10 min. After that, the volume was adjusted to 100 mL, then the acidified samples were centrifuged at 8,000 rpm for 10 min and filtered with 0.22 μm syringe water filters. For compounds chirality detection, the samples prepared above were further diluted 4-fold.

## Results

### Design and construction of D-lactic acid synthesis module in *G. thermoglucosidasius*

*G. thermoglucosidasius* DSM 2542 (wild type) contains the native *ldh* gene encoding L-lactate dehydrogenase, which produces L-lactic acid but not D-lactic acid (Fig. 5a). Therefore, to construct a D-lactic acid-producing strain, it is necessary to knock out *ldh* and introduce the *D-ldh* gene, which encodes D-lactate dehydrogenase. *G. thermoglucosidasius* is a high-temperature producer; therefore, the required enzymes should have good thermostability and catalytic performance. *D-ldh* from *B. licheniformis* BJQX combines these advantages. Therefore, gene *ldh* was deleted in the GTD0 strain, and the *D-ldh* gene was introduced at the same position under the control of the P*_ldh_* promoter. The P*_ldh_* promoter is relatively strong in *G. thermoglucosidasius*^23^. The obtained engineered strain GTD1 was verified by fed-batch fermentation. We found that it produced 37.5 g L^−1^ of D-lactic acid with an overall yield and productivity of 52.4% and 0.8 g L^−1^ h^−1^, respectively, within 48 h, (Fig. 2a), and few L-lactic acid was detected. The engineered strain GTD1 has the ability to produce D-lactic acid; however, we found that strain GTD1 still has many defects, including: a) slower glucose utilization rate in the middle and late stages of fermentation; b) formation of many by-products, mainly ethanol, which affects the yield; c) rapid death and lysis of large numbers of bacterial cells in the middle and late stages of fermentation, which is reflected by a rapid decrease in cell density; and d) intolerance to high concentrations of glucose (Fig. S1). Next, we individually solved these problems to improve the fermentation performance of the engineered strains.

**Fig. 2.**
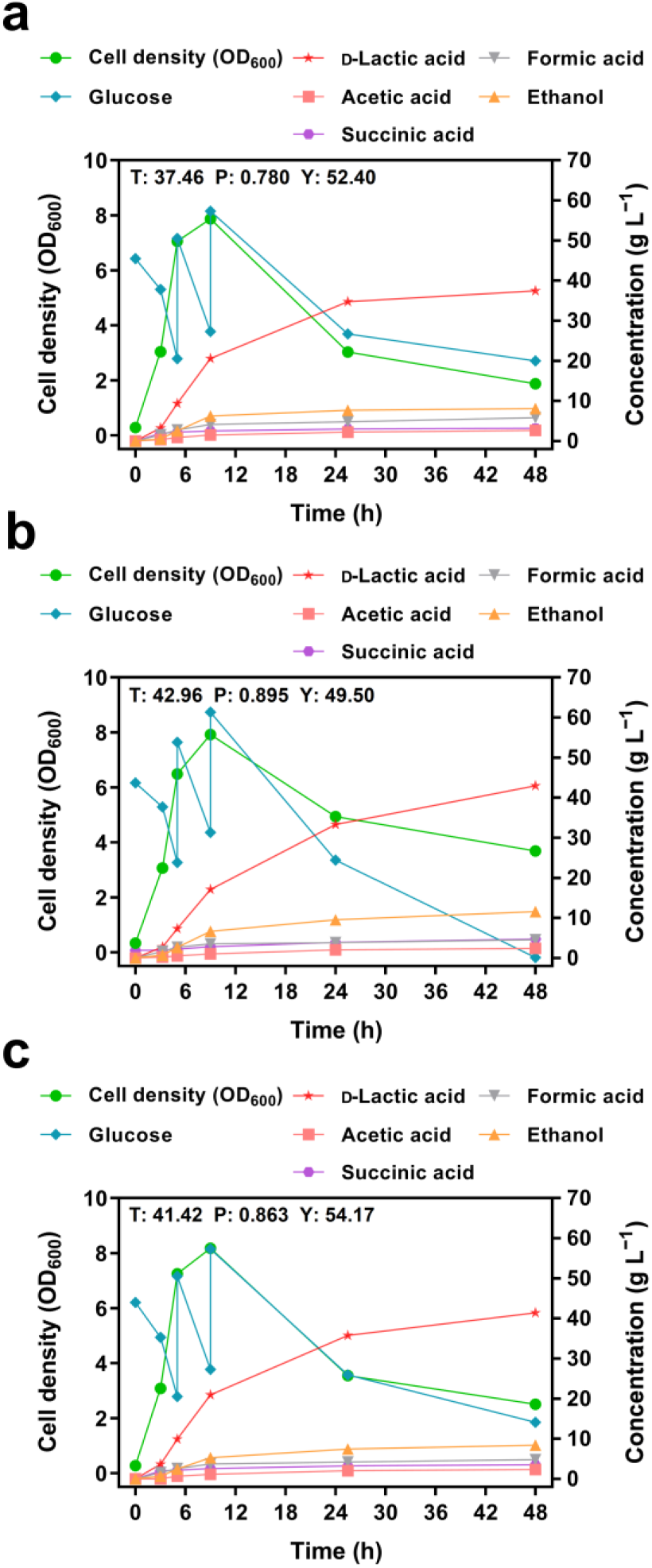
Fed-batch fermentations of engineered strains. Fed-batch fermentations of the **a** GTD1, **b** GTD2, and **c** GTD3 strains in a 5 L bioreactor with fermentation medium at 60°C, 80 rpm, and pH 7.0 (which was automatically maintained by Ca(OH)_2_), and the fermentation period was controlled within 50 h. The initial glucose concentration was 40 g L^−1^. T, P, and Y represent the titer (g L^−1^), productivity (g L^− 1^h^−1^), and yield (%, yields are based on per gram of fermentation product produced from per gram of glucose consumption) of the final sample, respectively. Symbols: green circle, cell density; blue diamond, glucose; red pentagram, D-lactic acid; grey inverted triangle, formic acid; pink square, acetic acid; orange triangle, ethanol; purple hexagon, succinic acid.

Pyruvate is generated from glucose during glycolysis and then catalyzed by D-lactate dehydrogenase to generate D-lactic acid (Fig. 1). Two key rate-limiting enzymes in glycolysis are 6-phosphofructokinase (encoded by *pfk*) and pyruvate kinase (encoded by *pyk*)^46^. Therefore, we introduced a copy of these genes into the engineered strain GTD1 to enhance the expression of these two enzymes. *B. coagulans* H-2 is a high-L-lactic acid-producing strain^4^, and its glycolytic pathway is relatively strong. Therefore, this was chosen to amplify *pfk* and *pyk*. Furthermore, we found that *pfk* and *pyk* were under the control of one promoter in their genome. We selected the P*_ldh_* promoter from the *ldh* gene in *G. stearothermophilus* and the P*_als_* promoter from the *alsS* gene in *B. licheniformis* BJQX, which are strong constitutive promoters, to control the expression of the *pfk* and *pyk*^2, 47^. We inserted these two gene expression cassettes into the engineered strain GTD1 to obtain strains GTD2 and GTD3. After verification of fed-batch fermentation (Fig. 2b, c), we found that both the engineered strains, GTD2 and GTD3, had improved glucose utilization rates in the middle and late stages of fermentation compared with strain GTD1 (Fig. 2a). The engineered strain GTD2 had the fastest glucose utilization rate in the middle and late stages of fermentation and consumed all the glucose within 48 h (the total mass of glucose added in the three fed-batch fermentations was equal). Therefore, the engineered strain GTD2 was used as the initial strain for subsequent genetic manipulation.

### Deletion of by-product (ethanol) synthesis in strain GTD2

Although the glucose utilization rate of strain GTD2 was improved in the middle and late stages of fermentation, its D-lactic acid titer and yield were not sufficiently high (Fig. 2b). We found that strain GTD2 produced a large amount of the following by- products during fermentation: 11.6 g L^−1^ ethanol, 4.8 g L^−1^ succinic acid, 4.6 g L^−1^ formic acid, and 2.4 g L^−1^ acetic acid. Since the fermentation temperature is 60°C, a large amount of ethanol volatilizes during the fermentation process. Hence, the actual ethanol titer is higher than the detected titer. Therefore, ethanol is the main by-product of the engineered strain GTD2. Acetaldehyde dehydrogenase catalyzes the generation of acetaldehyde from acetyl-CoA, which is then converted to ethanol by alcohol dehydrogenase (Fig. 1). Since acetaldehyde is toxic to cells, we intended to remove acetaldehyde dehydrogenase to prevent ethanol production. In addition, *G. thermoglucosidasius* contains an aldehyde-alcohol dehydrogenase (bifunctional dehydrogenase)^48^, which can directly catalyze acetyl-CoA to generate ethanol (Fig. 1); this was also a target of our deletion. We sequentially deleted *acdh1*, *acdh2*, and *aadh* from the engineered strain GTD2 to obtain GTD4, GTD5, and GTD6, respectively.

After verification through fed-batch fermentation (Fig. 3), we found that the growth status of the strain gradually deteriorated with the deletion of the ethanol pathway. This phenomenon is most evident in the engineered strain GTD6. The highest cell density of this strain was 5.9, which was much lower than that of the other engineered strains. Aldehyde dehydrogenase, alcohol dehydrogenase, aldehyde-alcohol dehydrogenase, and D-lactate dehydrogenase use NADH as a cofactor to catalyze related reactions. With the deletion of dehydrogenases, the NADH utilization pathways of the strain were reduced. Moreover, the D-lactate dehydrogenase expression in the current strain was not sufficiently strong. Consequently, the NADH utilization is limited, which in turn limits glycolysis. Therefore, the cells do not have sufficient energy for growth, which reduces the density of the strain. We solved this problem later in this paper. Furthermore, we found that the pathway catalyzed by aldehyde-alcohol dehydrogenase was the main ethanol synthesis pathway in this strain, which was reflected by an extremely low ethanol titer (0.7 g L^−1^) in the fermentation products of strain GTD6 (Fig. 3c). The engineered strain GTD6 produced 57.4 g L^−1^ of D-lactic acid with an overall yield and productivity of 76.8% and 1.1 g L^−1^ h^−1^, respectively (Fig. 3c), showing improved fermentation performance. Therefore, we selected this strain as the initial strain for subsequent genetic manipulation.

**Fig. 3.**
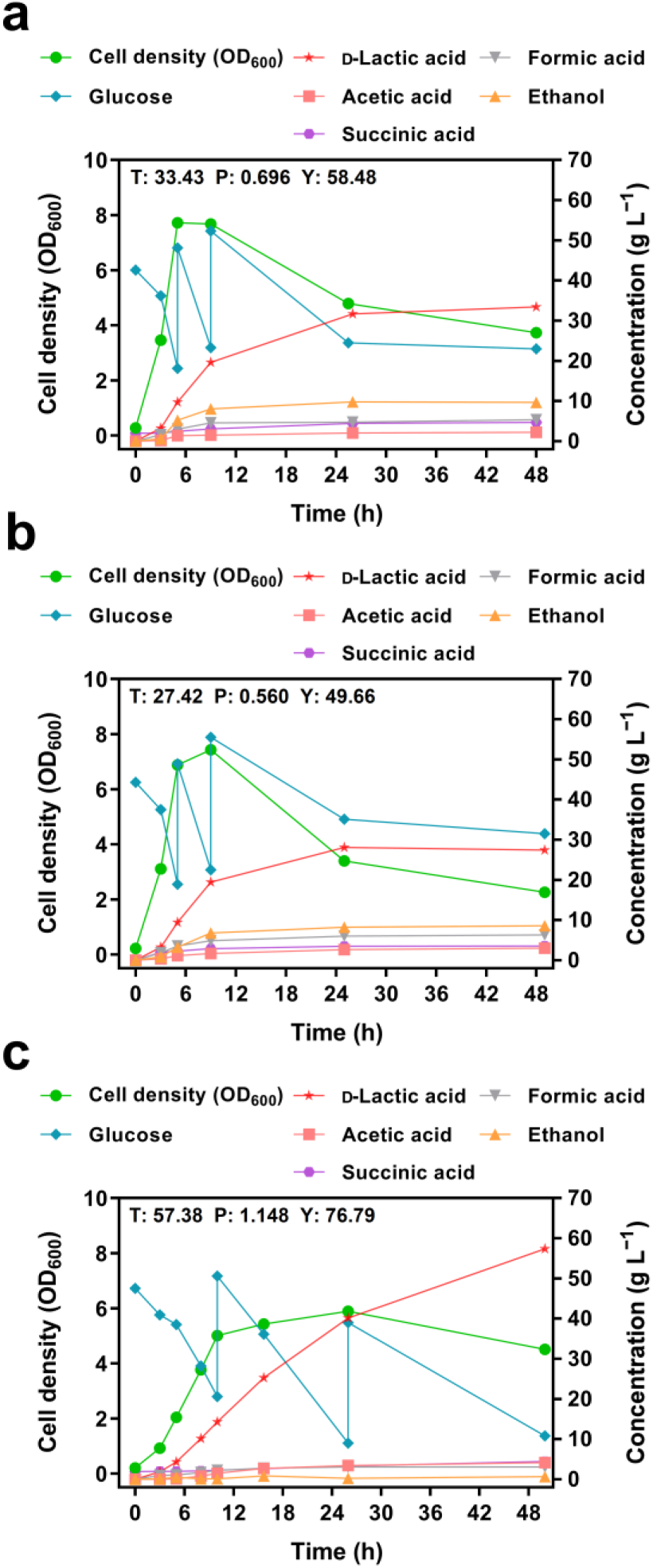
Fed-batch fermentations of engineered strains. Fed-batch fermentations of the **a** GTD4, **b** GTD5, and **c** GTD6 strains in a 5 L bioreactor with fermentation medium at 60°C, 80 rpm, and pH 7.0 (which was automatically maintained by Ca(OH)_2_), and the fermentation period was controlled within 50 h. The initial glucose concentration was 40 g L^−1^. T, P, and Y represent the titer (g L^−1^), productivity (g L^−1^ h^−1^), and yield (%, yields are based on per gram of fermentation product produced from per gram of glucose consumption) of the final sample, respectively. Symbols: green circle, cell density; blue diamond, glucose; red pentagram, D-lactic acid; grey inverted triangle, formic acid; pink square, acetic acid; orange triangle, ethanol; purple hexagon, succinic acid.

### Enhancement of D-lactic acid synthesis in strain GTD6

To solve the problem of poor growth in the engineered strain GTD6, we introduced another codon-optimized the D-*ldh* gene (Fig. 1). This improved the utilization of NADH in the strain, thereby promoting the progress of glycolysis. Moreover, the production capacity of D-lactic acid was further improved. We introduced the codon-optimized D-*ldh* gene at the position of the *aadh* gene in strain GTD6 to obtain the engineered strain GTD7. The codon-optimized D-*ldh* is under the control of the P*_aadh_* promoter, which is a strong promoter. After the verification using fed-batch fermentation (Fig. 4a), the cell growth status of the engineered strain GTD7 was restored and the production capacity of D-lactic acid was significantly improved (Fig. 3 and Fig. 4a). The engineered strain GTD7 produced 86.1 g L^−1^ of D-lactic acid with an overall yield and productivity of 92.1% and 1.8 g L^−1^ h^−1^, respectively (Fig. 4a).

**Fig. 4.**
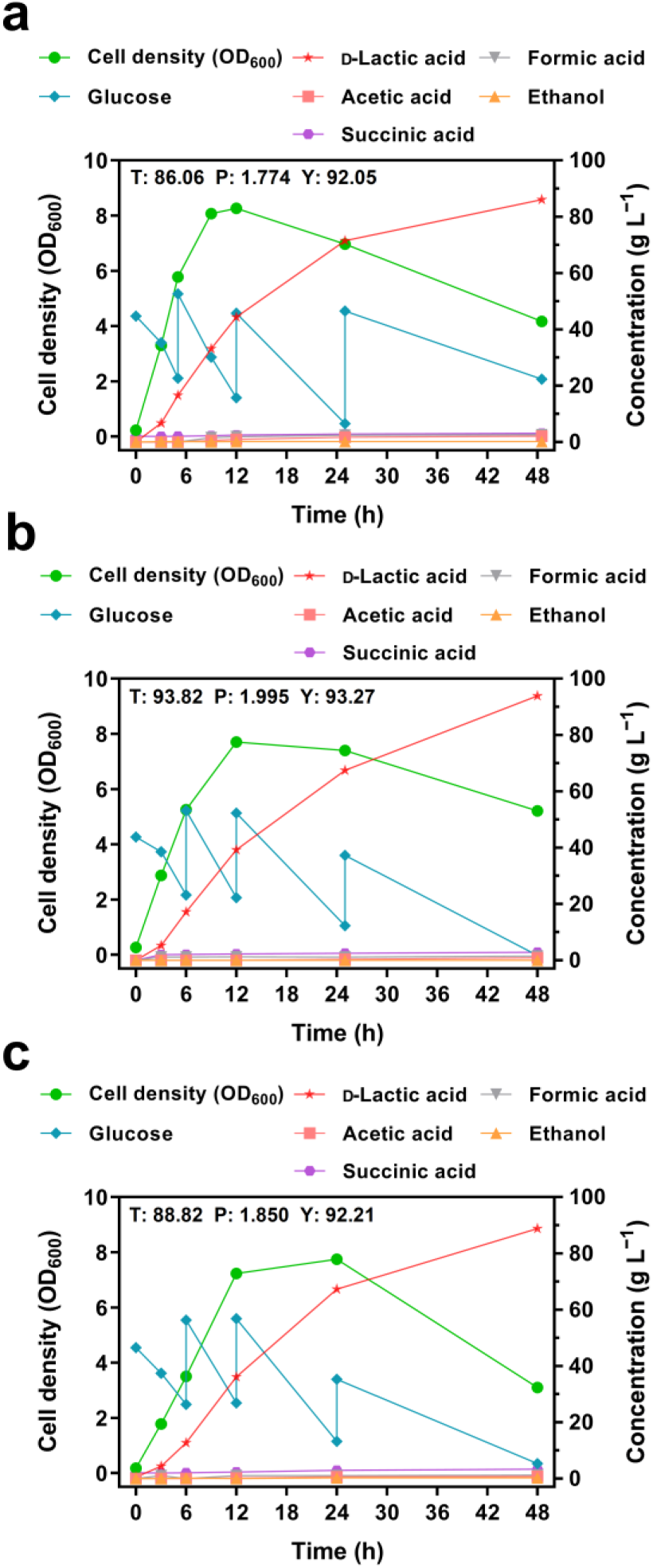
Fed-batch fermentations of engineered strains. Fed-batch fermentations of the **a** GTD7, **b** GTD8, and **c** GTD9 strains in a 5 L bioreactor with fermentation medium at 60°C, 80 rpm, and pH 7.0 (which was automatically maintained by Ca(OH)_2_), and the fermentation period was controlled within 50 h. The initial glucose concentration was 40 g L^−1^. T, P, and Y represent the titer (g L^−1^), productivity (g L^−1^ h^−1^), and yield (%, yields are based on per gram of fermentation product produced from per gram of glucose consumption) of the final sample, respectively. Symbols: green circle, cell density; blue diamond, glucose; red pentagram, D-lactic acid; grey inverted triangle, formic acid; pink square, acetic acid; orange triangle, ethanol; purple hexagon, succinic acid.

The yield reached an acceptable level; however, the titer and productivity of D-lactic acid required improvement. Furthermore, we tested the ability of the engineered strain GTD7 to utilize xylose, which produced 30.8 g L^−1^ of D-lactic acid with an overall yield and productivity of 88.3% and 0.6 g L^−1^ h^−1^, respectively, when using xylose as the carbon source (Fig. S2). This shows that stran GTD7 still has room for improvement when using xylose as a carbon source.

We found that the fermentation product of GTD7 contained 2.5 g L^−1^ formic acid and 2.1 g L^−1^ acetic acid (Fig. 4a and Table S3). Therefore, we removed the formate acetyltransferase and phosphate acetyltransferase to partially prevent the production of formic acid and acetic acid (Fig. 1). To further improve the D-lactic acid production capacity, we introduced an additional codon-optimized D-*ldh* gene. Therefore, we deleted *pflB* from the engineered strain GTD7 and introduced the codon-optimized *D-ldh* gene at the same position under the control of the P*_pflB_* promoter to obtain the engineered strain GTD8. We deleted *pta* from the engineered strain GTD8 to obtain GTD9. After fed-batch fermentation verifications (Fig. 4b, c), we found that the titers of formic acid and acetic acid did dropped to 1.4 g L^−1^ and 0.7 g L^−1^ in the engineered strains GTD8 and GTD9, respectively. This indicates that *G. thermoglucosidasius* has other formic and acetic acid production pathways. In addition, we found that the growth of the engineered strains GTD8 and GTD9 was slightly delayed compared to that of GTD7 (Fig. 4), indicating that deletion of the formic acid and acetic acid pathways had an effect on bacterial growth. Moreover, in the engineered strains GTD8 and GTD9, the titer of D-lactic acid did not increase significantly, and were 93.8 g L^−1^ and 88.8 g L^−1^, respectively. It was concluded that introduction of a copy of the codon-optimized D-*ldh* gene and deletion of *pflB* and *pta* genes did not significantly improve D-lactic acid production capacity. Therefore, we used GTD7 as the initial strain for subsequent genetic manipulation after comprehensive consideration.

### Construction and enhancement of L-lactic acid production in *G. thermoglucosidasius*

We also aimed to obtain a high-L-lactic acid-producing strain. The wild-type strain (GTD0) *G. thermoglucosidasius* DSM 2542 produces L-lactic acid. The strain GTD0 can produce 66.0 g L^−1^ of L-lactic acid with an overall yield and productivity of 78.4% and 1.4 g L^−1^ h^−1^, respectively (Fig. 5a). To improve the L-lactic acid production capacity of strain GTD0, we deleted the D-*ldh* gene, introduced the original *ldh* gene at the same position, and deleted the codon-optimized D-*ldh* gene in the engineered strain GTD7 to obtain the engineered strain GTD15. After the verification by fed-batch fermentation (Fig. 5b), we found that the production capacity of L-lactic acid was significantly improved in the engineered strain GTD15.

It produced 92.3 g L^−1^ of L-lactic acid with an overall yield and productivity of 95.3% and 1.9 g L^−1^ h^−1^, respectively, and the optical purity of 99.5% (Fig. 5b). We introduced an additional *ldh* gene to enhance the L-lactic acid synthesis. We introduced the original *ldh* gene and the *ldh* gene derived from *B. coagulans* H-2^4^ at the position of the *aadh* gene in the engineered strain GTD15 to obtain strains GTD16 and GTD17, respectively. After verification by fed-batch fermentation (Fig. 5c, d), we found that the L-lactic acid production capacity of the engineered strain GTD16 decreased significantly, which may be due to the presence of two identical *ldh* genes. During bacterial growth, these two identical genes may undergo homologous recombination that interferes with bacterial growth and L-lactic acid production. The engineered strain GTD17 produced 94.2 g L^−1^ of L-lactic acid with an overall yield and productivity of 91.5% and 2.0 g L^−1^ h^−1^, respectively, and an optical purity of 99.5% (Fig. 5d). The L-lactic acid production capacity of strain GTD17 was not significantly improved compared to that of GTD15. This indicates that the expression of L-lactate dehydrogenase in the engineered strain GTD15 is sufficient to meet the existing production capacity of the strain. Therefore, the L-lactic acid production capacity should be improved from other aspects.

### Exploring solutions for programmed cell death and lysis in *G. thermoglucosidasius*

Verification of the engineered strains using fed-batch fermentation (Fig. 2, 3, and 4) shows that a large number of bacterial cells died and lysed rapidly in the middle and late stages of fermentation, which was reflected by a rapid decrease in cell density. It has been reported that the toxin-antitoxin (TA) systems play a role in programmed cell death and lysis^49, 50^. We combined data from genome analysis and the TASmania database^51^ to identify two TA systems in the genome of *G. thermoglucosidasius* DSM 2542: the *ndoAI*/*ndoA* module and *mazE*/*mazF* module (Fig. 6a). Gene deletion experiments were used to explore whether these two TA systems could alleviate programmed cell death and lysis. We deleted the toxin genes *ndoA* and *mazF* and modules *ndoAI*/*ndoA*, and *mazE*/*mazF* in the engineered strain GTD7 to obtain strains GTD10, GTD11, GTD13, and GTD14, respectively. We deleted the toxin gene *mazF* from the engineered strain GTD10 to obtain GTD12. After verification by fed-batch fermentation (Fig. 6), we found that the D-lactic acid production capacity decreased in all engineered strains, especially in strains GTD10, GTD12, and GTD13.

Furthermore, programmed cell death and lysis were not alleviated in any engineered strain (Fig. 6). We also found that bacterial growth was significantly inhibited in the engineered strains GTD10, GTD12, and GTD13 (Fig. 6b, d, e). Therefore, we concluded that these two TA systems had a positive effect on the D-lactic acid production capacity and cell growth of *G. thermoglucosidasius* and that the *ndoAI*/*ndoA* module had a greater effect than the *mazE*/*mazF* module.

## Discussion

*G. thermoglucosidasius* is a new thermophilic chassis with excellent properties that deserves extensive research. *G. thermoglucosidasius* has been used to produce promising compounds at elevated temperatures, such as bioethanol^18, 47, 52–55^, isobutanol^56^, 2,3-butanediol^57^, terpenes^32^, and riboflavin^22, 58^. However, the production of these compounds is limited to the laboratory and cannot be scaled up for industrial production, mainly because of low product titers. The D-lactic acid production ability of the reprogrammed strain GTD7-144 in this study adheres to industrial requirement; this strain can produce 153.1 g L^−1^ of D-lactic acid with an overall yield and productivity of 93.0% and 3.2 g L^−1^ h^−1^ (Fig. 7b and Fig. 8), respectively, with an optical purity of 99.6% (Table S4). This also revealed the production potential of this thermophilic chassis.

This study promotes the industrial production of polymer-grade lactic acid. Using thermophilic microorganisms for fermentation at elevated temperatures significantly reduces production costs for fermentation at elevated temperatures^24, 59^. At present, the fermentation temperature of L-lactic acid- and D-lactic acid-producing strains is between 30 and 55°C ^2–4, 60^. In this study, the fermentation temperature (60°C) of *G. thermoglucosidasius* used to produce lactic acid was extremely high, and was higher than that used previously. This greatly reduces production costs. In addition, the GTP fermentation medium used in this study had a relatively low cost, which also reduced production costs. Furthermore, the strain GTD7-144 produced 153.1 g L^−1^ of D-lactic acid with an overall yield and productivity of 93.0% and 3.2 g L^−1^ h^−1^ (Fig. 7b and Fig. 8), respectively, with an optical purity of 99.6% (Table S4). The excellent D-lactic acid production performance also contributes to the industrialization of this strain. The engineered strains GTD15 and GTD17 also had good L-lactic acid production performance (Fig. 5 and Fig. 8), and will conform to industrial standards for L-lactic acid production following adaptive evolution.

Combination of rational and semi-rational design strategies is an important method for reprogramming chassis cells. We obtained optically pure lactic acid-producing strains GTD7 and GTD17 using the following series of rational metabolic engineering strategies: construction of the lactic acid synthesis module, deletion of by- product synthesis pathways, and enhancement of lactic acid synthesis pathway. However, rationally designed strains still have the following problems: lactic acid production capacity is not high enough, adaptability to barren medium is not strong enough, and growth state during the middle and late stages of fermentation is poor. Therefore, we performed semi-rational adaptive evolution to improve lactic acid production capacity and adaptability to barren media and identified relevant potential genetic targets. Programmed cell death and lysis are the main reasons for the poor growth of engineered strains during the middle and late stages of fermentation^49, 50^. The mechanisms underlying programmed cell death and lysis are complex and difficult to study^50^. This is due to DNA damage responses in some bacteria^61–63^. Kolodkin-Gal *et al.* reported that a linear pentapeptide is a quorumsensing factor required for *mazEF*-mediated cell death in *E. coli*^49^. Stein *et al.* reported that ageing exacerbates ribosome pausing to disrupt co-translational proteostasis, leading to a shortened cellular lifespan^64^. Researchers previously observed the same phenomenon in *G. thermoglucosidasius*^56^. In this study, we attempted to use a rational design strategy to solve this problem; however, we observed that the *ndoAI*/*ndoA* and *mazE*/*mazF* modules were not responsible for programmed cell death or lysis in this strain. Ultimately, we alleviated the problem of programmed cell death and lysis in the engineered strain by semi-rational adaptive evolution and found that genetic variants of genes related to transcriptional regulation, proteolysis, and stress response may be potential genetic targets. Semi-rational adaptive evolution solves these problems, which are intractable by rational design. Genome resequencing has also revealed potential genetic targets, all of which are worthy of further investigation for better rational designs.

To develop a thermophilic chassis *G. thermoglucosidasius* for the industrial production of promising compounds, many tasks must be performed. Although *G. thermoglucosidasius* has a wide spectrum of substrate utilization and catabolic versatility, its ability to utilize xylose needs to be improved when compared to its ability to utilize glucose (Fig. 4a and Fig. S2). Bashir *et al.* engineered *G. thermoglucosidasius* to produce bioethanol from wheat straw holocellulose^47^. Wang *et al.* explored the utilization of lignocellulose hydrolysates by *G. thermoglucosidasius*^58^. At present, more studies are needed to improve the utilization of inexpensive carbon sources. Meanwhile, enabling technologies suitable for this strain also need to be developed to better serve this chassis, including shorter cycle gene manipulation methods, more comprehensive promoter and RBS libraries, mining of enzymes with good thermostability and catalytic performance, and more accurate metabolic model predictions. Furthermore, researchers need to broaden the types of compounds produced by *G. thermoglucosidasius* to improve its production value.

In summary, we obtained optically pure L-lactic acid- and D-lactic acid-producing strains, GTD17 and GTD7, respectively, by modifying *G. thermoglucosidasius* using rational metabolic engineering strategies (Fig. 8). We then carried out semi-rational adaptive evolution of the engineered strain GTD7 to further improve its D-lactic acid production performance. The final strain GTD7-144 produced 153.1 g L^−1^ of D-lactic acid with an overall yield and productivity of 93.0% and 3.2 g L^−1^ h^−1^, respectively. The optical purity of D-lactic acid was 99.6%. Additionally, genome resequencing and analysis of GTD7-144 revealed potential genetic targets that could improve the fermentation performance of the strain. This study reveals the production potential of *G. thermoglucosidasius* at the quasi-industrial level and represents an advancement in the industrial production of polymer-grade lactic acid.

## Competing interests

The authors declare that they have no competing interests.

## Acknowledgements

This study is supported by the grant from National Natural Science Foundation of China (22138007 and 32170105). We thank Dr. Zilong Li (Institute of Microbiology, Chinese Academy of Sciences) for providing the plasmid pUB-sfGFP.

## Notes

### Competing Interest Statement

The authors have declared no competing interest.

## References

1 Garvie, E. I. Bacterial lactate dehydrogenases. Microbiol. Rev. 44, 106–139, doi:10.1128/mr.44.1.106–139.1980 (1980).

2 Li, C., Tao, F. & Xu, P. Carbon flux trapping: highly efficient production of polymer-grade D-lactic acid with a thermophilic D-lactate dehydrogenase. Chembiochem 17, 1491–1494, doi:10.1002/cbic.201600288 (2016).

3 Han, X. et al. Steps toward high-performance PLA: economical production of D-lactate enabled by a newly isolated *Sporolactobacillus terrae* strain. Biotechnol. J. 14, e1800656, doi:10.1002/biot.201800656 (2019).

4 Zhang, F. et al. Kinetic characteristics of long-term repeated fed-batch (LtRFb) L-lactic acid fermentation by a *Bacillus coagulans* strain. Eng. Life Sci. 20, 562–570, doi:10.1002/elsc.202000043 (2020).

5 Tan, C., Tao, F. & Xu, P. Direct carbon capture for the production of high-performance biodegradable plastics by cyanobacterial cell factories. Green Chem. 24, 4470–4483, doi:10.1039/d1gc04188f (2022).

6 Swetha, T. A. et al. A review on biodegradable polylactic acid (PLA) production from fermentative food waste - its applications and degradation. Int. J. Biol. Macromol. 234, 123703, doi:10.1016/j.ijbiomac.2023.123703 (2023).

7 Ikada, Y., Jamshidi, K., Tsuji, H. & Hyon, S. H. Stereocomplex formation between enantiomeric poly(lactides). Macromolecules 20, 904–906, doi:10.1021/ma00170a034 (1987).

8 Farah, S., Anderson, D. G. & Langer, R. Physical and mechanical properties of PLA, and their functions in widespread applications – a comprehensive review. Adv. Drug Deliv. Rev. 107, 367–392, doi:10.1016/j.addr.2016.06.012 (2016).

9 Fukushima, K., Chang, Y. H. & Kimura, Y. Enhanced stereocomplex formation of poly(L-lactic acid) and poly(D-lactic acid) in the presence of stereoblock poly(lactic acid). Macromol. Biosci. 7, 829–835, doi:10.1002/mabi.200700028 (2007).

10 Yu, B. et al. Morphology and internal structure control over PLA microspheres by compounding PLLA and PDLA and effects on drug release behavior. Colloids Surf. B Biointerfaces 172, 105–112, doi:10.1016/j.colsurfb.2018.08.037 (2018).

11 Purnama, P., Samsuri, M. & Iswaldi, I. Properties enhancement of high molecular weight polylactide using stereocomplex polylactide as a nucleating agent. Polymers (Basel) 13, 1725, doi:10.3390/polym13111725 (2021).

12 Jalali, A., Romero-Diez, S., Nofar, M. & Park, C. B. Entirely environment-friendly polylactide composites with outstanding heat resistance and superior mechanical performance fabricated by spunbond technology: exploring the role of nanofibrillated stereocomplex polylactide crystals. Int. J. Biol. Macromol. 193, 2210–2220, doi:10.1016/j.ijbiomac.2021.11.052 (2021).

13 Nanda, S., Patra, B. R., Patel, R., Bakos, J. & Dalai, A. K. Innovations in applications and prospects of bioplastics and biopolymers: a review. Environ. Chem. Lett. 20, 379–395, doi:10.1007/s10311-021-01334-4 (2022).

14 Nazina, T. N. et al. Taxonomic study of aerobic thermophilic bacilli: descriptions of *Geobacillus subterraneus* gen. nov., sp. nov. and *Geobacillus* uzenensis sp. nov. from petroleum reservoirs and transfer of Bacillus stearothermophilus, Bacillus thermocatenulatus, Bacillus thermoleovorans, Bacillus kaustophilus, Bacillus thermoglucosidasius and Bacillus thermodenitrificans to Geobacillus as the new combinations G. stearothermophilus, G. thermocatenulatus, G. thermoleovorans, G. kaustophilus, G. thermoglucosidasius and G. thermodenitrificans. Int. J. Syst. Evol. Microbiol. 51, 433–446, doi:https://doi.org/10.1099/00207713-51-2-433 (2001).

15 Hussein, A. H., Lisowska, B. K. & Leak, D. J. The genus *Geobacillus* and their biotechnological potential. Adv. Appl. Microbiol. 92, 1–48, doi:10.1016/bs.aambs.2015.03.001 (2015).

16 Liang, J., Roberts, A., van Kranenburg, R., Bolhuis, A. & Leak, D. J. Relaxed control of sugar utilization in *Parageobacillus thermoglucosidasius* DSM 2542. Microbiol. Res. 256, 126957, doi:10.1016/j.micres.2021.126957 (2022).

17 Mol, V. et al. Genome-scale metabolic modeling of *P. thermoglucosidasius* NCIMB 11955 reveals metabolic bottlenecks in anaerobic metabolism. Metab. Eng. 65, 123–134, doi:10.1016/j.ymben.2021.03.002 (2021).

18 Cripps, R. E. et al. Metabolic engineering of *Geobacillus thermoglucosidasius* for high yield ethanol production. Metab. Eng. 11, 398–408, doi:10.1016/j.ymben.2009.08.005 (2009).

19 Bacon, L. F., Hamley-Bennett, C., Danson, M. J. & Leak, D. J. Development of an efficient technique for gene deletion and allelic exchange in *Geobacillus* spp. Microb. Cell. Fact. 16, 58, doi:10.1186/s12934-017-0670-4 (2017).

20 Sheng, L., Kovacs, K., Winzer, K., Zhang, Y. & Minton, N. P. Development and implementation of rapid metabolic engineering tools for chemical and fuel production in *Geobacillus thermoglucosidasius* NCIMB 11955. Biotechnol. Biofuels 10, 5, doi:10.1186/s13068-016-0692-x (2017).

21 Frenzel, E., Legebeke, J., van Stralen, A., van Kranenburg, R. & Kuipers, O. P. In vivo selection of sfGFP variants with improved and reliable functionality in industrially important thermophilic bacteria. Biotechnol. Biofuels 11, 8, doi:10.1186/s13068-017-1008-5 (2018).

22 Yang, Z. et al. Engineering thermophilic *Geobacillus thermoglucosidasius* for riboflavin production. Microb. Biotechnol. 14, 363–373, doi:10.1111/1751-7915.13543 (2021).

23 Lau, M. S. H., Sheng, L., Zhang, Y. & Minton, N. P. Development of a suite of tools for genome editing in *Parageobacillus thermoglucosidasius* and their use to identify the potential of a native plasmid in the generation of stable engineered strains. ACS Synth. Biol. 10, 1739–1749, doi:10.1021/acssynbio.1c00138 (2021).

24 Abdel-Banat, B. M., Hoshida, H., Ano, A., Nonklang, S. & Akada, R. High-temperature fermentation: how can processes for ethanol production at high temperatures become superior to the traditional process using mesophilic yeast? Appl. Microbiol. Biotechnol. 85, 861–867, doi:10.1007/s00253-009-2248-5 (2010).

25 Zeldes, B. M. et al. Extremely thermophilic microorganisms as metabolic engineering platforms for production of fuels and industrial chemicals. Front. Microbiol. 6, 1209, doi:10.3389/fmicb.2015.01209 (2015).

26 Wu, Y. et al. Eliminating host-guest incompatibility via enzyme mining enables the high-temperature production of *N*-acetylglucosamine. iScience 26, 105774, doi:10.1016/j.isci.2022.105774 (2023).

27 Choi, K. R. et al. Systems metabolic engineering strategies: integrating systems and synthetic biology with metabolic engineering. Trends Biotechnol. 37, 817–837, doi:10.1016/j.tibtech.2019.01.003 (2019).

28 Bhalla, A., Bansal, N., Kumar, S., Bischoff, K. M. & Sani, R. K. Improved lignocellulose conversion to biofuels with thermophilic bacteria and thermostable enzymes. Bioresour. Technol. 128, 751–759, doi:10.1016/j.biortech.2012.10.145 (2013).

29 Robak, K. & Balcerek, M. Review of second generation bioethanol production from residual biomass. Food Technol. Biotechnol. 56, 174–187, doi:10.17113/ftb.56.02.18.5428 (2018).

30 Kruger, A., Schafers, C., Schroder, C. & Antranikian, G. Towards a sustainable biobased industry - highlighting the impact of extremophiles. N. Biotechnol. 40, 144–153, doi:10.1016/j.nbt.2017.05.002 (2018).

31 Losa, J. et al. Perspective: a stirring role for metabolism in cells. Mol. Syst. Biol. 18, e10822, doi:10.15252/msb.202110822 (2022).

32 Styles, M. Q. et al. The heterologous production of terpenes by the thermophile *Parageobacillus thermoglucosidasius* in a consolidated bioprocess using waste bread. Metab. Eng. 65, 146–155, doi:10.1016/j.ymben.2020.11.005 (2021).

33 Wu, Y., Jameel, A., Xing, X. H. & Zhang, C. Advanced strategies and tools to facilitate and streamline microbial adaptive laboratory evolution. Trends Biotechnol. 40, 38–59, doi:10.1016/j.tibtech.2021.04.002 (2022).

34 Ling, C. et al. Muconic acid production from glucose and xylose in *Pseudomonas putida* via evolution and metabolic engineering. Nat. Commun. 13, 4925, doi:10.1038/s41467-022-32296-y (2022).

35 Kim, K. et al. Adaptive laboratory evolution of *Escherichia coli* W enhances gamma-aminobutyric acid production using glycerol as the carbon source. Metab. Eng. 69, 59–72, doi:10.1016/j.ymben.2021.11.004 (2022).

36 Keller, P. et al. Generation of an *Escherichia coli* strain growing on methanol via the ribulose monophosphate cycle. Nat. Commun. 13, 5243, doi:10.1038/s41467-022-32744-9 (2022).

37 Pereira, R. et al. Adaptive laboratory evolution of tolerance to dicarboxylic acids in *Saccharomyces cerevisiae*. Metab. Eng. 56, 130–141, doi:10.1016/j.ymben.2019.09.008 (2019).

38 Matson, M. M. et al. Adaptive laboratory evolution for improved tolerance of isobutyl acetate in *Escherichia coli*. Metab. Eng. 69, 50–58, doi:10.1016/j.ymben.2021.11.002 (2022).

39 Gao, J., Li, Y., Yu, W. & Zhou, Y. J. Rescuing yeast from cell death enables overproduction of fatty acids from sole methanol. Nat. Metab. 4, 932–943, doi:10.1038/s42255-022-00601-0 (2022).

40 Caspeta, L. & Nielsen, J. Thermotolerant yeast strains adapted by laboratory evolution show trade-off at ancestral temperatures and preadaptation to other stresses. mBio 6, e00431, doi:10.1128/mBio.00431-15 (2015).

41 Pereira, R. et al. Elucidating aromatic acid tolerance at low pH in *Saccharomyces cerevisiae* using adaptive laboratory evolution. Proc. Natl. Acad. Sci. U. S. A. 117, 27954–27961, doi:10.1073/pnas.2013044117 (2020).

42 Tian, X. et al. Metabolic engineering coupled with adaptive evolution strategies for the efficient production of high-quality L-lactic acid by *Lactobacillus paracasei*. Bioresour. Technol. 323, 124549, doi:10.1016/j.biortech.2020.124549 (2021).

43 Horton, R. M., Hunt, H. D., Ho, S. N., Pullen, J. K. & Pease, L. R. Engineering hybrid genes without the use of restriction enzymes: gene splicing by overlap extension. Gene 77, 61–68, doi:https://doi.org/10.1016/0378-1119(89)90359-4 (1989).

44 Zhao, B. et al. Kinetics of D-lactic acid production by *Sporolactobacillus* sp. strain CASD using repeated batch fermentation. Bioresour. Technol. 101, 6499–6505, doi:10.1016/j.biortech.2010.03.069 (2010).

45 Jiang, S., Xu, P. & Tao, F. L-Lactic acid production by *Bacillus coagulans* through simultaneous saccharification and fermentation of lignocellulosic corncob residue. Bioresour. Technol. Rep. 6, 131–137, doi:10.1016/j.biteb.2019.02.005 (2019).

46 Zhao, C., Lin, Z., Dong, H., Zhang, Y. & Li, Y. Reexamination of the physiological role of PykA in *Escherichia coli* revealed that it negatively regulates the intracellular ATP levels under anaerobic conditions. Appl. Environ. Microbiol. 83, e00316–00317, doi:10.1128/AEM.00316-17 (2017).

47 Bashir, Z. et al. Engineering *Geobacillus thermoglucosidasius* for direct utilisation of holocellulose from wheat straw. Biotechnol. Biofuels 12, 199, doi:10.1186/s13068-019-1540-6 (2019).

48 Extance, J., Danson, M. J. & Crennell, S. J. Structure of an acetylating aldehyde dehydrogenase from the thermophilic ethanologen *Geobacillus thermoglucosidasius*. Protein Sci. 25, 2045–2053, doi:10.1002/pro.3027 (2016).

49 Kolodkin-Gal, I., Hazan, R., Gaathon, A., Carmeli, S. & Engelberg-Kulka, H. A linear pentapeptide is a quorum-sensing factor required for *mazEF*-mediated cell death in *Escherichia coli*. Science 318, 652–655, doi:10.1126/science.1147248 (2007).

50 Allocati, N., Masulli, M., Di Ilio, C. & De Laurenzi, V. Die for the community: an overview of programmed cell death in bacteria. Cell Death Dis. 6, e1609, doi:10.1038/cddis.2014.570 (2015).

51 Akarsu, H. et al. TASmania: a bacterial Toxin-Antitoxin Systems database. PLoS Comput. Biol. 15, e1006946, doi:10.1371/journal.pcbi.1006946 (2019).

52 Van Zyl, L. J., Taylor, M. P., Eley, K., Tuffin, M. & Cowan, D. A. Engineering pyruvate decarboxylase-mediated ethanol production in the thermophilic host *Geobacillus thermoglucosidasius*. Appl. Microbiol. Biotechnol. 98, 1247– 1259, doi:10.1007/s00253-013-5380-1 (2014).

53 Zhou, J., Wu, K. & Rao, C. V. Evolutionary engineering of *Geobacillus thermoglucosidasius* for improved ethanol production. Biotechnol. Bioeng. 113, 2156–2167, doi:10.1002/bit.25983 (2016).

54 Raita, M., Ibenegbu, C., Champreda, V. & Leak, D. J. Production of ethanol by thermophilic oligosaccharide utilising *Geobacillus thermoglucosidasius* TM242 using palm kernel cake as a renewable feedstock. Biomass Bioenergy 95, 45–54, doi:10.1016/j.biombioe.2016.08.015 (2016).

55 Calverley, J., Zimmerman, W. B., Leak, D. J. & Hemaka Bandulasena, H. C. Continuous removal of ethanol from dilute ethanol-water mixtures using hot microbubbles. Chem. Eng. J. 424, 130511, doi:10.1016/j.cej.2021.130511 (2021).

56 Lin, P. P. et al. Isobutanol production at elevated temperatures in thermophilic *Geobacillus thermoglucosidasius*. Metab. Eng. 24, 1–8, doi:10.1016/j.ymben.2014.03.006 (2014).

57 Zhou, J., Lian, J. & Rao, C. V. Metabolic engineering of Parageobacillus thermoglucosidasius for the efficient production of (2R, 3R)-butanediol. Appl. Microbiol. Biotechnol. 104, 4303–4311, doi:10.1007/s00253-020-10553-8 (2020).

58 Wang, J. et al. Dynamic control strategy to produce riboflavin with lignocellulose hydrolysate in the thermophile *Geobacillus thermoglucosidasius*. ACS Synth. Biol. 11, 2163–2174, doi:10.1021/acssynbio.2c00087 (2022).

59 Yang, X. Manual of Industrial Microbiology and Biotechnology Ch. 47 (ASM Press, Washington, 2010).

60 Juturu, V. & Wu, J. C. Microbial production of lactic acid: the latest development. Crit. Rev. Biotechnol. 36, 967–977, doi:10.3109/07388551.2015.1066305 (2015).

61 Bejerano-Sagie, M. et al. A checkpoint protein that scans the chromosome for damage at the start of sporulation in *Bacillus subtilis*. Cell 125, 679–690, doi:10.1016/j.cell.2006.03.039 (2006).

62 Witte, G., Hartung, S., Buttner, K. & Hopfner, K. P. Structural biochemistry of a bacterial checkpoint protein reveals diadenylate cyclase activity regulated by DNA recombination intermediates. Mol. Cell 30, 167–178, doi:10.1016/j.molcel.2008.02.020 (2008).

63 Kreuzer, K. N. DNA damage responses in prokaryotes: regulating gene expression, modulating growth patterns, and manipulating replication forks. Cold Spring Harb. Perspect. Biol. 5, a012674, doi:10.1101/cshperspect.a012674 (2013).

64 Stein, K. C., Morales-Polanco, F., van der Lienden, J., Rainbolt, T. K. & Frydman, J. Ageing exacerbates ribosome pausing to disrupt cotranslational proteostasis. Nature 601, 637–642, doi:10.1038/s41586-021-04295-4 (2022).

